# B cells transiently exposed to antigen that return to quiescence respond to protein immunization similarly to naive B cells

**DOI:** 10.1101/303420

**Authors:** Jackson S. Turner, Fang Ke, Irina Grigorova

## Abstract

Transient exposure of B cells to moderately multivalent antigens has been shown to lead to their activation and recruitment into T-dependent B cell response in the presence of T cell help. However, when T cell help is delayed, transiently activated B cells return to a naïve-like state. While B cells that briefly encountered cognate antigen and then underwent inactivation are not tolerant, whether they may be hypo-or hyper-responsive to cognate antigen is not known. In this study we show that naïve B cells and those previously exposed to antigen and then inactivated generate comparable germinal center and plasma cell responses after immunization with a wide range of antigen concentrations.

## Introduction

Induction of T-dependent B cell response is essential for mounting long-term high affinity humoral (antibody) immunity against pathogens and is the goal for most existing vaccines. For recruitment into the T-dependent response, B cells need to acquire foreign antigen (Ag) via cognate B cell receptor (BCR), which triggers initial activation of B cells (signal I). This leads to internalization of Ag, its proteolysis in the endosomes and antigenic peptides loading onto Major Histocompatibility class II Molecules (MHCII) followed by presentation of MHCII/peptide complexes on the plasma membrane for recognition by cognate Ag-specific helper T cells (Th). The second signal provided by Th cells promotes sustained B cell activation and triggers their proliferation with subsequent differentiation into germinal center (GC) B cells and antibody-secreting short-lived plasma cells (1–4).

Different factors may affect recruitment of various Ag-specific B cell clones into T-dependent responses (5–7). One of these factors is the duration of B cell exposure to Ag and the timing of T cell help. Continuous exposure of B cells to Ag in the absence of T cell help is known to induce B cell tolerance through either anergy or activation-induced cell death (AICD) (8–10). However, the duration of B cell exposure to foreign Ags may vary depending on the Ag size, biophysical properties, preexisting Abs and other factors (11). Large Ags have been shown to be initially restricted to the subcapsular, medulary, and cortical sinuses, as well as the interfollicular areas of Ag-draining lymph nodes (12–15). Based on data from 2-photon intravital microscopy the initial exposure of B cells to the Ags at these sites is likely to be transient (16, 17). In a recent study we showed that transient exposure to cognate Ag induces B cell activation that is sufficient for B cell recruitment into the immune response when T cell help is available (18). However, if T cell help is delayed, in the absence of recurrent BCR stimulation, B cells return to a naïve-like state. They downregulate activation markers, reduce presentation of foreign antigenic peptides, and become unresponsive to T cell help within 48 h. We also showed that in contrast to B cells that reacquire Ags for prolonged period of time, B cells transiently exposed to monovalent or moderately multivalent antigens do not undergo AICD *in vivo.* Finally, these inactivated B cells do not become tolerant, as they responded to immunization with cognate Ag by mounting GC and plasmablasts (PB, precursors of plasma cells) responses. Moreover, when Ag and T cell help are available in excess, inactivated B cells generate similar early GC and PB responses to those of naïve Ag-specific cells (18). However, whether B cells that have returned to quiescence have similar responsiveness as naïve B cells to Ag and T cell help when they are not saturating or whether their sensitivity is modified by their previous exposure to Ag has not been assessed.

In the current study we compared the ability of B cells transiently exposed to cognate Ag and then inactivated with naïve B cells to mount immune responses after immunization with various doses of Ag *in vivo*. Our findings suggest that naïve and inactivated B cells have similar responsiveness to the cognate protein Ag.

## Materials and Methods

### Mice

C57BL/6 (B6) mice were purchased from Charles River Laboratories and Ptprc^a^ Pepc^b^/BoyJ (B6-CD45.1) mice were purchased from the Jackson Laboratory. MD4 BCR transgenic (Ig-Tg) mice (B6 background) (19) were generously provided by Jason Cyster. MD4 mice were crossed with BoyJ (B6-CD45.1) mice and maintained on this background. Donor and recipient mice were 6-12 weeks of age. All mice were maintained in a specific pathogen free environment and protocols were approved by the Institutional Animal Care and Use Committee of the University of Michigan.

### Ag preparation

Ovalbumin (OVA) was purchased from Sigma. Duck eggs were locally purchased and duck egg lysozyme (DEL) was purified and conjugated to OVA via glutaraldehyde cross-linking as previously described (20).

### Immunization and adoptive transfer

MD4 Ig-Tg B cells were enriched from male and female donor mice by negative selection as previously described (21). For transient exposure to Ag, purified MD4 B cells were incubated with 2.5 μg/mL DEL-OVA *ex vivo* for 5 minutes at 37 °C, washed four times with room temperature DMEM supplemented with 4.5 g/L glucose, L-glutamine and sodium pyruvate, 2% FBS, 10 mM HEPES, 50 IU/mL of penicillin, and 50 mg/mL of streptomycin. 5×10^3^ of DEL-OVA-pulsed or naïve MD4 B cells were then transferred i.v. to recipient mice. At 7 days after the cell transfer male recipient mice were immunized i.p. with 50, 5 or 0.5 mg DEL-OVA in Ribi adjuvant (Sigma), prepared according to the manufacturer’s directions.

### Flow cytometery

The following antibodies specific to B220 (RA3-6B2), CD95 (Jo2), from BD-Pharmingen; CD45.1 (A20), CD3 (17A2), from Biolegend; CD38 (90), GL-7 (GL-7), from eBioscience have been used for flow cytometry analysis. Single-cell suspensions of splenocytes were incubated with fluorophore-conjugated antibodies for 20 minutes on ice and washed twice with 200 μl FACS buffer (2% FBS, 1mM EDTA, 0.1% NaN_3_ in PBS). For intracellular staining, surface-stained cells were fixed and permeabilized for 20 minutes on ice with BD Cytofix/Cytoperm buffer, washed twice with 200 μl BD Perm/Wash buffer, incubated with Alexa 647-conjugated Hen Egg Lysozyme (which has very high affinity to MD4 B cells’ BCR) for 30 minutes on ice, followed by two more washes with 200 μl Perm/Wash buffer, and resuspended in FACS buffer for acquisition. Cells were acquired on a FACSCanto, and data was analyzed using FlowJo (TreeStar).

### Statistics

Statistical tests were performed as indicated using Prism 7 (GraphPad). Differences between groups not annotated by an asterisk did not reach statistical significance. No blinding or randomization was performed for animal experiments, and no animals or samples were excluded from analysis.

## Results and Discussion

In the previous study we showed that transient exposure of MD4 Ig-Tg B cells (with BCRs specific to Duck Egg Lysozyme, DEL) to 0.5-50 μg/mL of moderately multivalent DEL-OVA (DEL conjugated to ovalbumine) Ag *ex vivo* led to their robust activation and recruitment into the GC and PB response *in vivo* when T cell help was immediately available (18). However, in the absence of T cell help, Ag exposed Ig-Tg B cells returned to naïve-like inactivated state within 2-3 days of their residence within secondary lymphoid organs. When transferred into mice preimmunized with OVA and reimmunized with 50 μg of DEL-OVA, which is expected to provide an excess of Ag and T cell help, the naïve or inactivated Ig-Tg B cells developed similar responses.

In the current study we have modified our previously utilized experimental scheme to make the activation conditions more limiting. Ig-Tg B cells were pulsed *ex vivo* with 2.5 μg/mL DEL-OVA for 5 min at 37° C and then extensively washed. 5×10^3^ of naïve or DEL-OVA pulsed Ig-Tg B cells were then transferred into unimmunized recipient mice. Seven days later, after inactivation of Ig-Tg B cells, the recipient mice were immunized with increasing doses of DEL-OVA in Ribi adjuvant. Seven days later we assessed Ig-Tg B cells’ recruitment into the GC and PB responses by flow cytometry analysis (Fig. 1A, B).

**Figure 1.**
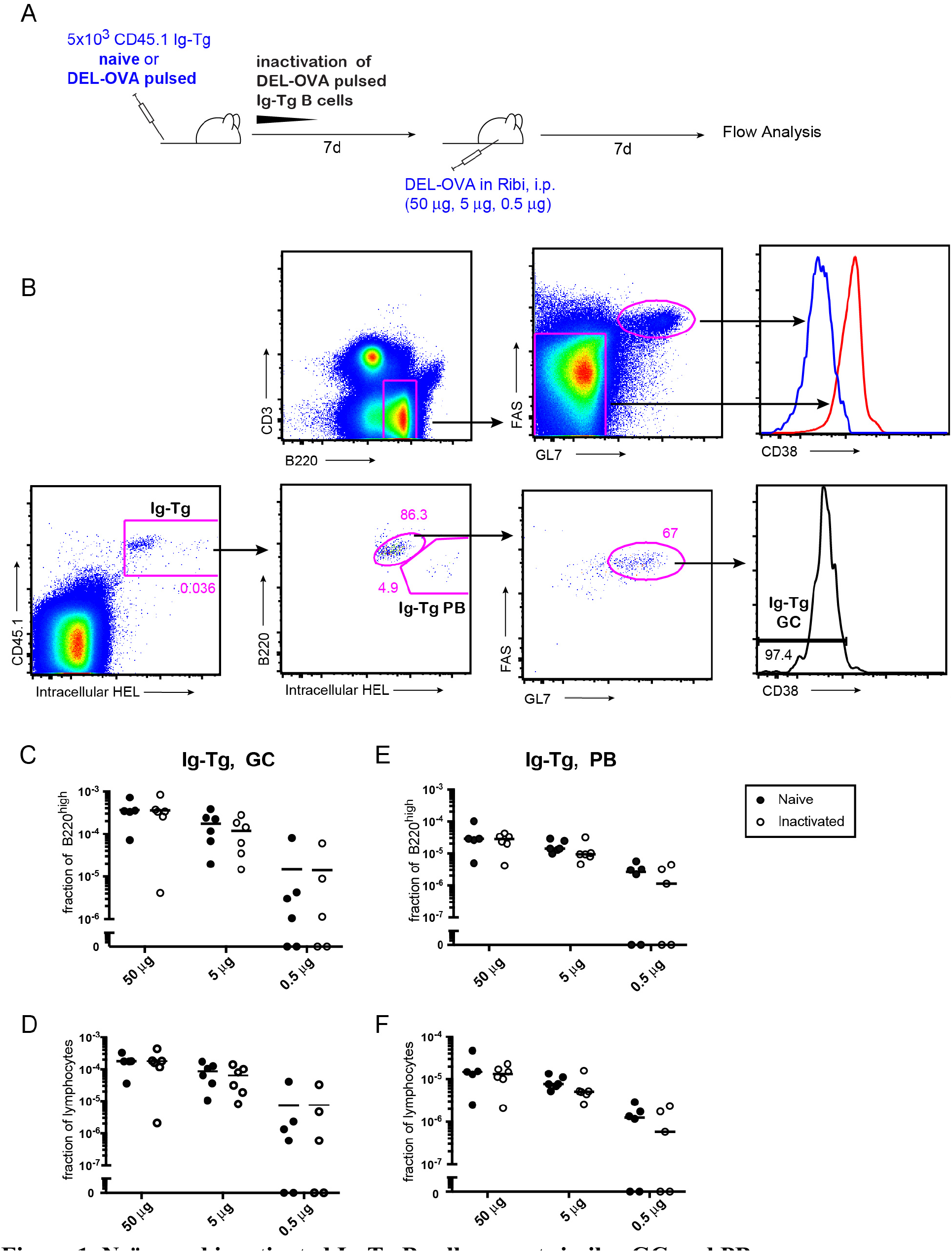
Naïve and inactivated Ig-Tg B cells mount similar GC and PB response after immunization with various doses of DEL-OVA Ag. **A**, Experimental outline. Purified MD4 Ig-Tg B cells were pulsed *ex vivo* for 5 min with 2.5 μg/mL DEL-OVA and then extensively washed. 5×10^3^ Ag-pulsed or naïve B cells were transferred into recipient B6 mice. 7 days later the recipients were immunized i.p. with indicated doses of DEL-OVA in Ribi adjuvant. **B**, The flow cytometry gating strategy to identify Ig-Tg GC B cells (CD45.1^+^ HEL^interm^. B220^high^ FAS^high^ GL7^high^ CD38^low^) and PB (CD45.1^+^ HEL^high^ B220^low^). Gated on lymphocytes. **C-F**, Ig-Tg GC (C, D) and PB (E,F) cell frequency in respect to B220^high^ (C, E) and total (D, F) splenocytes at 7 days following immunization. Data from 3 independent experiments, 1-2 mice per condition. Each symbol represents one mouse. Lines represent mean values. No statistical significant differences between naïve and inactivated GC B cells was identified by T-test.

Immunization of mice with a very low dose of DEL-OVA Ag (0.5 μg/mL) led to little or no recruitment of Ig-Tg B cells into the GC and PB response; their participation progressively increased after administration of intermediate (5 μg/mL) and high doses (50 μg/mL) of Ag (Fig. 1 C-F). In all cases, independently of the dose of the Ag utilized, naïve or inactivated Ig-Tg B cells had similar participation in the GC and PB response (Fig. 1C-F). Of note, as previously demonstrated, when fewer than 10^4^ Ig-Tg B cells are transferred per mouse, the Ig-Tg GC and PB responses to DEL-OVA are not saturated (18). These results suggest that Ig-Tg B cells that underwent Ag-dependent activation and then returned to quiescence in the absence of T cell help have similar sensitivity to cognate protein Ags as naïve Ig-Tg B cells in their ability to engage into the T-dependent response upon provision of cognate Ag and T cell help.

Overall, based on our previous studies and current work we suggest that in the absence of T cell help, transient exposure of B cells to moderately multivalent Ags is not sufficient to induce B cell AICD or anergy, or to cause hypo or hyper-responsiveness to protein Ag and T cell help (18).

